# Bacterial community dynamics during embryonic development of the little skate (*Leucoraja erinacea*)

**DOI:** 10.1101/2020.12.30.424594

**Authors:** Katelyn Mika, Alexander S. Okamoto, Neil H. Shubin, David B. Mark Welch

## Abstract

**Background:** Microbial transmission from parent to offspring is hypothesized to be widespread in vertebrates. However, evidence for this is limited as many evolutionarily important clades remain unexamined. There is currently no data on the microbiota associated with any Chondrichthyan species during embryonic development, despite the global distribution, ecological importance, and phylogenetic position of this clade. In this study, we take the first steps towards filling this gap by investigating the microbiota associated with embryonic development in the little skate, *Leucoraja erinacea*, a common North Atlantic species and popular system for chondrichthyan biology.

**Methods:** To assess the potential for bacterial transmission in an oviparous chondrichthyan, we used 16S rRNA amplicon sequencing to characterize the microbial communities associated with the skin, gill, and egg capsule of the little skate, at six points during ontogeny. Community composition was analyzed using the QIIME2 pipeline and microbial continuity between stages was tracked using FEAST.

**Results:** We identify site-specific and stage-specific microbiota dominated by the bacterial phyla *Proteobacteria* and *Bacteroidetes*. This composition is similar to, but distinct from, that of previously published data on the adult microbiota of other chondrichthyan species. Our data reveal that the skate egg capsule harbors a highly diverse bacterial community–particularly on the internal surface of the capsule–and facilitates intergenerational microbial transfer to the offspring. Embryonic skin and external gill tissues host similar bacterial communities; the skin and gill communities later diverge as the internal gills and skin denticles develop.

**Conclusions:** Our study is the first exploration of the chondrichthyan microbiota throughout ontogeny and provides the first evidence of vertical transmission in this group.

## Introduction

Host-associated microbial communities are often species and tissue-specific due to complex local interactions between hosts and microbes [1, 2]. Species can acquire their microbiota through three possible processes: horizontal microbial transmission between conspecifics, vertical microbial transmission from parents to offspring, or similar environmental sourcing across the host species [3]. In the first case, microbes can be horizontally transferred between conspecifics during social interactions and sexual behaviors, potentially homogenizing the bacterial communities across the host population. Alternatively, vertical transmission allows parents to directly provide their progeny with symbiotic microbial taxa [4]. Lastly, microbes can be recruited from the surrounding environment through the host’s contact with fluids, substrates, or diet, allowing for rapid changes in community composition in response to external conditions over the lifespan of an individual. The relative contributions of these transmission modes likely covary with life history, balancing the need for intergenerational continuity of genomic information with the capacity to respond to environmental cues [3, 5].

Next-generation sequencing has facilitated the characterization of a broad range of microbiomes across an increasing diversity of host species; nonetheless, many important marine clades remain understudied. Chondrichthyans—the earliest branching of the extant, jawed-vertebrate lineages —constitute one of the major divisions of vertebrates [6]. To date, only a limited number of culture-independent studies of chondrichthyan microbiota have been conducted [7–13]. Existing studies of chondrichthyans all focus on species belonging to subclass Elasmobranchii, which includes sharks, skates, rays, and guitarfish. This literature is skewed towards the skin or gut microbiota of pelagic sharks [7–11, 13] or the skin microbiota of select ray species [11, 12]. These datasets show that elasmobranch skin microbiota differ from that of the surrounding environment and are primarily dominated by the phyla *Proteobacteria* and *Bacteroidetes*, similar to the skin microbiota of other marine species [14–16]. However, this work is limited to adult elasmobranchs, providing no direct information on juvenile microbiota or intergenerational transmission in chondrichthyans.

In some clades, microbiome composition closely tracts host phylogeny over evolutionary time, resulting in long-term eco-evolutionary relationships known as phylosymbiosis [17]. Previous research has identified signatures of phylosymbiosis in elasmobranchs by showing a correlation between host phylogenetic distance and the taxonomic composition of the microbiota [11]. Of the three processes of transmission described, horizontal transmission has the most limited explanatory potential for this finding as chondrichthyan species are largely asocial with aggregations driven primarily by environmental factors or reproduction [18, 19]. Vertical transmission or environmental sourcing are more promising potential mechanisms to explain the signature of phylosymbiosis. Data on environmental sourcing in chondrichthyans are limited [9, 12] and are difficult to acquire, while vertical transmission has been unexplored. Given the lack of data and the hypothesis that vertical transmission is widespread in vertebrates [4], we investigated the potential for vertical transmission in the model chondrichthyan, the little skate (*Leucoraja erinacea*).

Oviparity is present in almost half of chondrichthyans and may be the plesiomorphic reproductive mode for this clade [6, 20]. Like other skates (family: Rajidae), little skates are egg-laying elasmobranchs that protect their embryos inside egg capsules, colloquially known as a mermaid’s purses [21]. Development of the egg capsule starts in the nidamental organ where the posterior half is formed before the fertilized egg is deposited into the capsule, at which point the capsule is rapidly sealed shut [22]. These capsules are then laid on the seafloor and the embryos develop inside for months to years depending upon the temperature [23, 24] and species [25]. While egg capsules can osmoregulate at all stages, they are initially sealed to anything larger than small molecules, e.g. glucose and urea can pass through but insulin cannot [26, 27]. Slits at the anterior and posterior ends of egg capsule open up late in development allowing seawater to flow through [28]. The potential effects of this environmental shift on the microbiota and host development are unknown. Upon hatching, juvenile skates are self-sufficient, with no known parental care [6]. These life history traits–long embryonic development and lack of parental care after oviposition–pose potential obstacles to vertical microbial transmission in members of this clade. Thus, skates are a useful system for testing vertical transmission because confounding parental contact is minimized and any transmitted microbial community is likely stable over a substantial period of time.

The little skate is a model system for research in chondrichthyan embryology and development [24, 29–32] with a sequenced genome [33]. This species is common in the North Atlantic [34] and easy to obtain through sampling and breeding methods implemented in that region. Little skate embryos have gestational periods inside the egg capsule of 22–54 weeks depending on the season [23]. Embryogenesis is divided into thirty-three stages, based on morphological features [35, 36]. In this study, our goal was to characterize the microbiota associated with embryonic development of the little skate and to assess the potential for vertical transmission in this species. To accomplish this, we used 16S rRNA amplicon sequencing to describe and track changes in bacterial diversity throughout little skate ontogeny by sampling the microbiota of the skin and gills, as well as the internal liquid and internal surface of the egg capsule at six developmental stages. These stages are (i) stage 0, when capsules are freshly laid and fertilization cannot be visually confirmed; (ii) stage 16, an early stage when the embryo can be visually identified; (iii) stage 26, by which external gill filaments have formed and the egg capsule is still sealed; (iv) stage 30, when the egg capsule starts to open and the gills remain external; (v) stage 33, by which time the egg capsule is open and the embryo is fully formed with internal gills; and (vi) adult. These stages span the duration of embryonic development from shortly after oviposition until just before hatching and capture distinct periods related to organogenesis and environmental exposure.

## Materials and Methods

### Sample Collection

Adult skates used in this study were wild caught in the Northern Atlantic and housed in 15°C filtered seawater (400-micron mesh followed by sand filtration) at the Marine Resources Center (MRC) of the Marine Biological Laboratory, Woods Hole, MA. All embryos were laid within the facility tanks by this adult population. Samples were collected from adult females (*n*=4) by separately swabbing the gills and the skin around the cloaca. Adult females were not directly associated with any particular egg capsule used in this study but were all sexually mature, housed in the MRC breeding tanks, and thus serve as representative, potential mothers. Egg capsules were sampled at five timepoints: stages 0, 16, 26, 30, and 33 (*n*=4 each, *n*=20 total) as per refs. [35, 36]. Capsules were windowed with a razor blade and the embryos euthanized by cervical transection. At each stage, samples of the internal liquid (*n*=20) were collected using a 1000 mL pipette and the inside of the egg capsule was swabbed (*n*=20), as shown in supplementary figure 1. All samples at stage 33 were open, as were two samples (A & D) at stage 30. All other samples were closed. Samples where the egg capsule slits were already open to the environment were drained into a collection tube before the egg capsule was windowed. At stages 26, 30 and 33, gill filament samples and tail clippings (∼2 cm long) were collected (*n*=4 each, *n=2*4 total). Control samples included hand swabs of A.S.O and K.M. and bench swabs before sample processing both on the day of sample collection and again on the day of DNA extraction (*n=* 6 total). A sample of the bench after sterilization (*n*=1) was taken on the day of sample collection as well. Sterile water was collected as a negative control. To broadly sample the marine bacteria likely to be encountered by skates in the MRC, 1 mL of water was collected in 1.5 mL tubes from (i) ambient-temperature water tanks (*n*=2); (ii) 15°C tanks housing the skates (*n*=2); (iii) ocean water from the dock neighboring the pump into the MRC (*n*=2); and (iv) the bucket used to transfer the skate embryos from the MRC to the dissection station was collected before (*n=*1) and after sampling (*n*=1). Prior to sample collection, the bench and all dissection tools were sterilized using Clorox bleach, followed by 70% ethanol. These surfaces were re-sterilized with 70% ethanol between each egg capsule and with bleach and ethanol between every four egg capsules. All skate samples were collected on the same day and egg capsules were opened in a randomly selected order.

The DNeasy PowerSoil Kit (Qiagen, Hilden Germany) was used for isolation of the microbial DNA from each sample. FLOQSwabs (COPAN, Murrieta CA) were used to collect all swab samples. Swabs trimmed to fit or tissue samples from the gills and tail were placed directly in the PowerSoil Kit PowerBead tubes after collection. For all liquid samples, 200 µL of the sample was added to the corresponding tube. After collection, samples were left at -20°C overnight prior to completion of the extraction protocol. Extraction continued according to the manufacturer’s instructions. The negative control and post-cleaning bench swab failed to amplify, suggesting our sterilization technique was effective (Supplementary Table S1). Pre-cleaning bench samples were thus unlikely to have contaminated other samples and were excluded from further analysis.

### Sequencing and Library Preparation

To identify the bacterial community within each sample, the V4-V5 region of the 16S gene was amplified and sequenced at the Keck Environmental Genomics Facility at the Marine Biological Laboratory as described [37]. Amplification was done using forward primer (518F) CCAGCAGCYGCGGTAAN and reverse primers (926R) CCGTCAATTCNTTTRAGT, CCGTCAATTTCTTTGAGT, and CCGTCTATTCCTTTGANT. Three reverse primers were used to capture known 16S sequence diversity more effectively than a single highly degenerate primer. PCR cycle structure was 94°C for 2 minutes, 30 cycles of repeating 94°C for 30° seconds, 57°C for 45 seconds, and 72°C for 1 minute, followed by 72°C for 2 minutes then a hold at 4°C. Sequencing was done on an Illumina MiSeq platform. Results were then uploaded to the Visualization and Analysis of Microbial Population Structures (VAMPS) website (https://vamps2.mbl.edu/) [38]. Raw data was obtained and deposited at the NCBI Short Read Archive (PRJNA688288).

Denoising to address amplicon errors, classifying to identify the taxonomic affiliation of each sequence, and alpha and beta diversity methods to assess community composition were all implemented using QIIME2 version 2019.10 [39]. Demultiplexed paired-end sequencing data was denoised without any trimming and chimeras removed using the DADA2 QIIME2 plugin [40]. The resulting amplicon sequence variants (ASVs) were classified by training a Naive Bayes classifier using the SILVA (132 release) 16S only, 99% identity clustered sequences [41–43]. ASVs were collapsed to a maximum specificity of seven taxonomic levels, which corresponds to the species level. Data were normalized using transform_sample_counts in Phyloseq [44, 45] by dividing the count per ASV within a sample by the total count for that sample, followed by multiplying this ratio by 10,000.

### Microbial Community Analysis

QIIME2 was used to calculate Pielou’s evenness, Shannon and Chao1 alpha diversity indices [46], and the Bray-Curtis dissimilarity index for beta diversity [47]. Beta diversity differences between sample types were visualized using principal coordinate analysis. Between-group significance levels for alpha and beta diversity were assessed using Kruskal-Wallis [48] and PERMANOVA [49] tests, respectively, with a Benjamini-Hochberg false discovery rate (FDR) correction. The significance threshold for these tests was set at q<0.05. Taxa comprising the common core microbiota [50] of the (i) egg capsule, (ii) combined external gill and embryonic skin, (iii) internal gill, and (iv) adult skin were identified using the *feature-table core-features* function in QIIME2 at a threshold of 75% presence. This threshold was chosen to prioritize taxa with high levels of occupancy in tissues of interest. Given our small sample size for each tissue and stage, thresholds higher than 75% result in very few core taxa, while lowering this threshold results in rapid increases in core taxa numbers.

LEfSe (Linear Discriminant Analysis (LDA) Effect Size) was used to identify significantly enriched taxa and implemented in the Galaxy web application (http://huttenhower.org/galaxy/) [51] using a *P*-value cut-off of 0.05, an LDA score cut-off of 2, and a one-against-all strategy. FEAST [52] in R v3.6.2 [53] was used to track bacterial community continuity between stages, with samples from the preceding stage (unless otherwise noted), water, and the investigators’ hands used as potential sources. R Studio (Version 1.2.5033) with the *ggplot2* package [54] was used to produce all figures.

## Results

Eighty-four out of eighty-eight samples successfully amplified and were sequenced to produce a total of 3,516,842 reads and 41,486 ASVs (Supplementary Tables S1 and S2). These ASVs were classified into 2,255 unique taxonomic identities using the SILVA database. Each sample contained between 170 and 147,651 reads, with a median value of 36,480. ASV assignments ranged from 1 to 33,211 reads, with a median value of 29. Rarefaction curves of ASVs recovered versus sequencing depth showed that sequencing depth was sufficient to discover the majority of ASVs in a sample (Supplementary Fig. 2).

### Taxonomic characterization of *L. erinacea* microbiota

Skate and egg capsule samples are dominated by ASVs of the phyla *Proteobacteria* (58% of ASVs), *Bacteroidetes* (21%), and *Planctomycetes* (5%). The dominant classes identified are *Gammaproteobacteria* (33%), *Alphaproteobacteria* (21%), *Bacteroidia* (21%), and *Planctomycetacea* (4%) (Fig. 1). Adult skin samples are uniquely enriched for *Bacteroidetes*, which accounts for 81% of the bacterial community, while in all other samples *Proteobacteria* are most abundant. Within the control samples, the water samples are dominated by *Proteobacteria* (87%), followed by *Bacteroidetes* (5%), *Cyanobacteria* (2.6%), and *Actinobacteria* (2%). The investigators’ hand samples are quite distinct from all other samples, dominated by the phyla *Actinobacteria* (49%), *Cyanobacteria* (31%), *Proteobacteria* (9%) and *Firmicutes* (5%), showing the characteristic microbial community skew associated with ocean water exposure [55]. As is typical when investigating poorly studied environments, fewer ASVs can be categorized at finer phylogenetic resolutions, with <98% being classified at the phylum level, 97% at the class level, <83% at the order level, <70% at the family level and <60% at the genus level. The most common families identified are *Rhodobacteraceae* (8.1%), *Flavobacteriaceae* (6.9%), *Enterobacteraceae* (6.1%), *Saprospiraceae* (3.2%), and *Devosiaceae* (3.2%) (Supplementary Fig. 3). The most common genera are *Escherichia-Shigella* (6.1%), *Cutibacterium* (3.2%), *Devosia* (2.0%), and *Lutibacter* (0.9%) (Supplementary Fig. 4).

**Figure 1:**
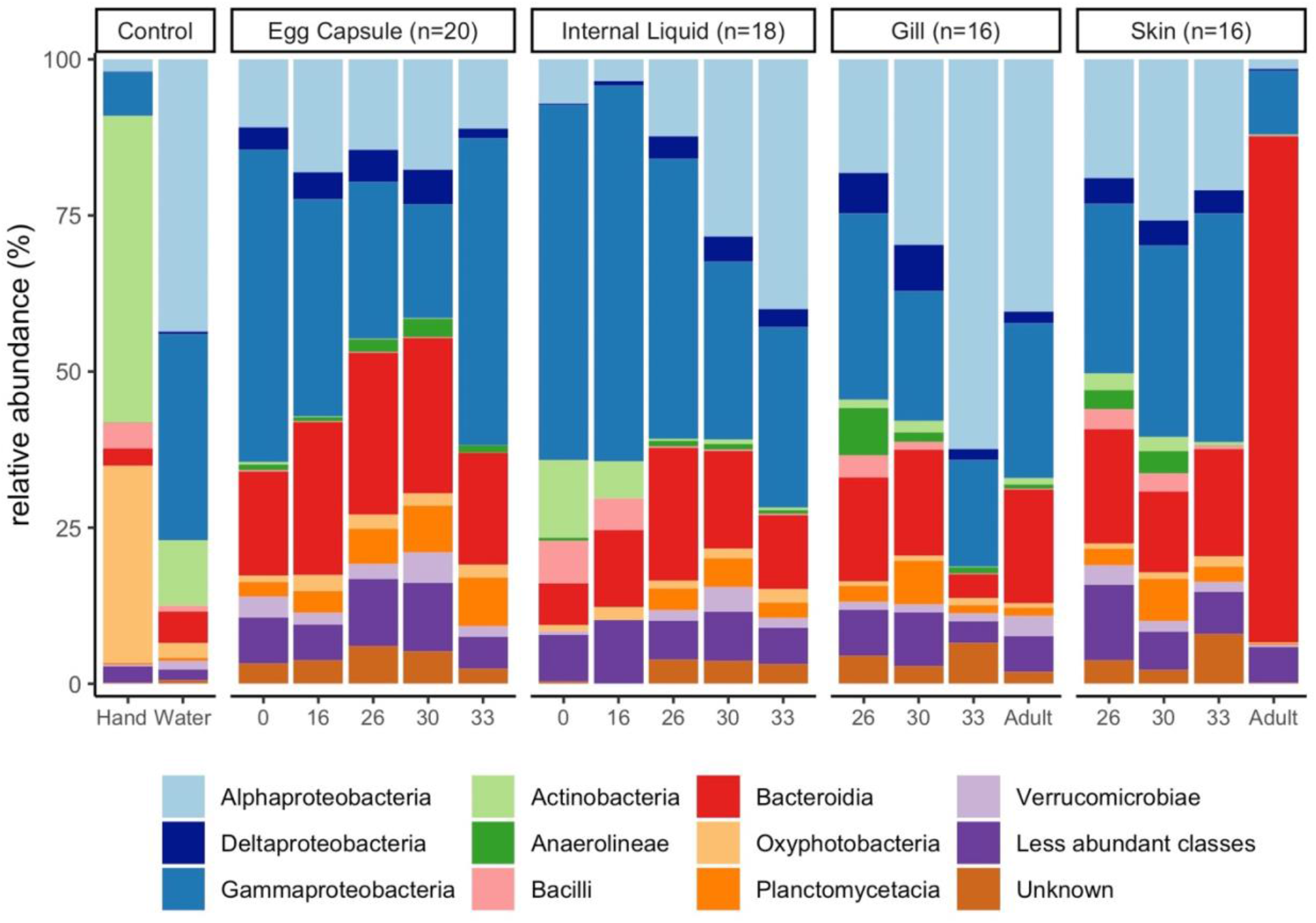
Taxonomic composition of embryonic and adult skate bacterial communities. Relative abundance of the top ten bacterial classes in the dataset are shown for each site and timepoint as well as for water and hand controls. First, classes of phylum *Proteobacteria* are shown in shades of blue, followed by other classes ordered alphabetically. For the controls, n=4 for hand and n=8 for water samples.

### Beta and alpha diversity of skate bacterial communities

We explored variation in microbial community composition between samples–Beta diversity–by sample tissue and stage using Bray-Curtis dissimilarity (Fig. 2). Due to the low number of internal liquid samples at stage 16 which successfully amplified (n=2), these samples cannot be statistically differentiated from other samples but cluster tightly with stage 0 internal liquid.Water is likewise distinct from all skate tissues (q<0.05) except internal liquid from stage 33 embryos (q<0.08), at which time the internal liquid contains a significant amount of water due to the egg capsule opening. All egg capsule samples form a tight cluster distinct (q<0.08) from all other samples except mid-stage internal liquid (stages 16-30; q<0.50). Gill tissue samples split into two clusters (q <0.05): the external, embryonic gill stages (26-30), and the internal, later stages (33-Adult). Internal gill samples are distinct from egg capsule (q<0.05) and late skin samples (stage 33-Adult, q<0.05) while external gill samples (stage 26-30) cannot be distinguished from early skin samples (stages 26-30; q>0.20). Stage 30, 33 and adult skin are all significantly different (q<0.05) from each other. Adult skin is distinct from all other samples (q<0.05) and separates from all others along PC3 (Fig. 2B). Samples at stage 30 show no clear separation on open or closed status for any of the tissues examined.

**Figure 2:**
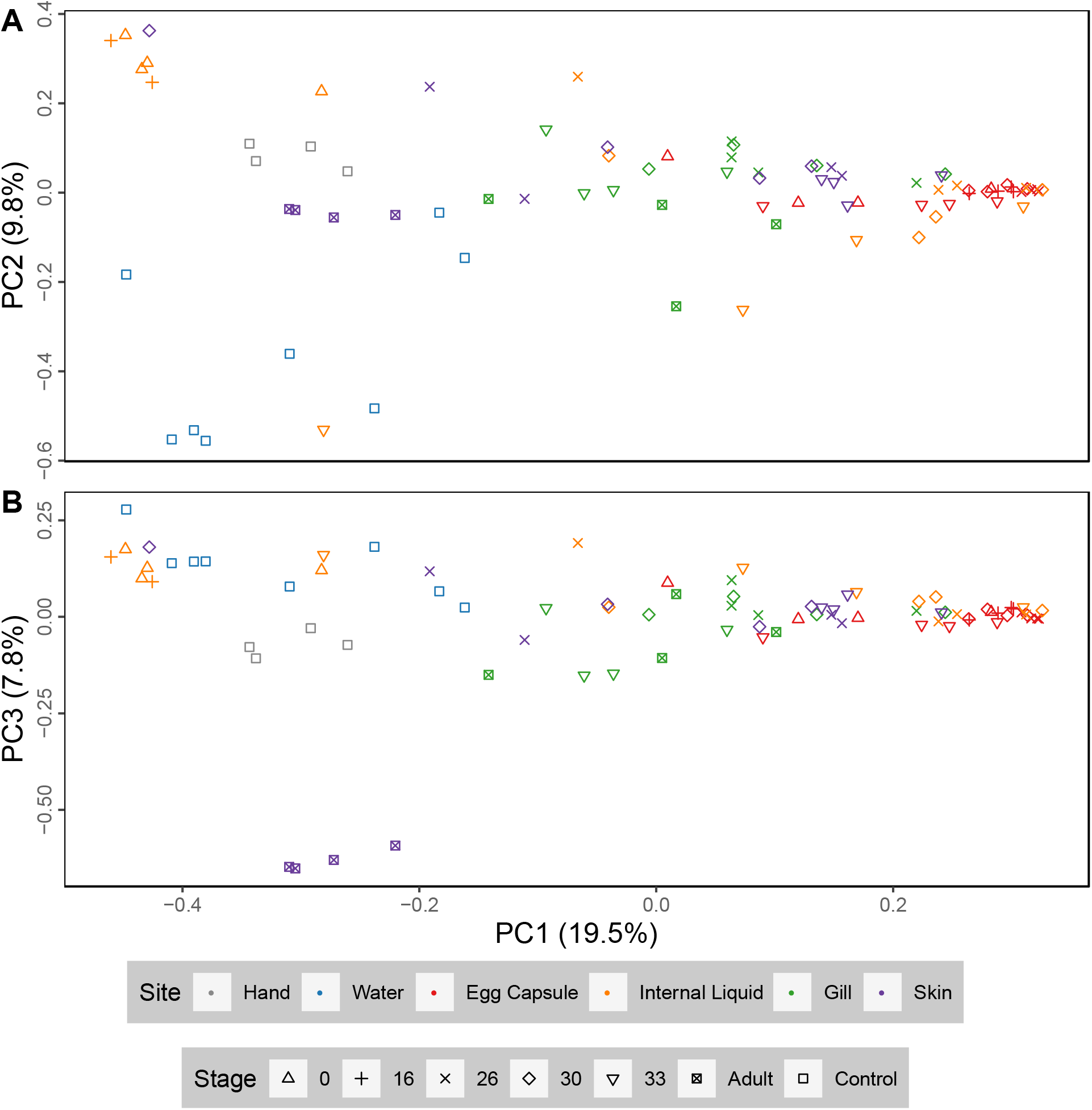
Principal coordinate analysis plots of little skate bacterial samples. PCoA analysis (Bray-Curtis) of skate and control samples. PC1 versus PC2 (A) and PC1 versus PC3 (B). Sample distribution the same as in Figure 1.

Principal coordinate axes generated using the full dataset are primarily driven by the differences between sampling locations, which may obscure tissue-specific clustering patterns. Therefore, we stratified the data into individual tissues and reran Bray-Curtis dissimilarity analyses on the reduced datasets (Supplementary Fig. 5). Egg capsule samples, which have indistinct subclustering in Fig. 2, show more substructure when analyzed independently, with stages 0, 30, and 33 forming unique subclusters (q<0.05; Supplementary Fig. 5A). Internal liquid has no statistically significant subclusters (q>0.09; Supplementary Fig. 5B). Bray-Curtis dissimilarity analysis on all gill samples supports the internal and external gill subclusters seen in the full dataset (Supplementary Fig. 5C; q<0.05). Adult skin is distinct from all other skin (Supplementary Fig. 5D; q<0.05) along PC1 which explains 36.5% of the variance within skin samples. Due to high variability in stage 26 and 30 skin, stage 33 skin is statistically differentiated (q<0.05) but clusters tightly with the majority of these samples.

We explored diversity within a sample (alpha diversity), in two different ways: we used the Shannon index to assess both observed richness (the observed number of ASVs present in each sample) and evenness (relative abundance of ASVs in each sample): and the Chao1 index to assess the estimated total richness of the tissue site (the number of ASVs in the population represented by each sample) (Fig. 3). Total ASV counts and Pielou’s evenness for each sample are listed in Supplementary Table S2. Shannon and Chao1 indices were calculated for each sample site, splitting gills into external (stages 26-30) and internal (stages 33-Adult), skin into embryonic (stages 26-33) and adult, and averaging across all stages for egg capsule and internal liquid. The total richness of the egg capsule is significantly greater than all other sampling sites by Chao1 (pairwise Kruskal-Wallis, q<0.05). The total richness of the internal liquid increases with stage, but this trend is not significant; a similar trend is not seen in the egg capsule. When considering observed richness and evenness, egg capsule samples have a significantly higher mean Shannon index than all other sites except external gills. Taken together this suggests that while external gills have a relatively small number of taxa, the community is much more evenly distributed than most other sites. In contrast, adult skin has a small number of taxa dominated by a few highly abundant taxa. No other pairwise comparisons were significant using either metric.

**Figure 3:**
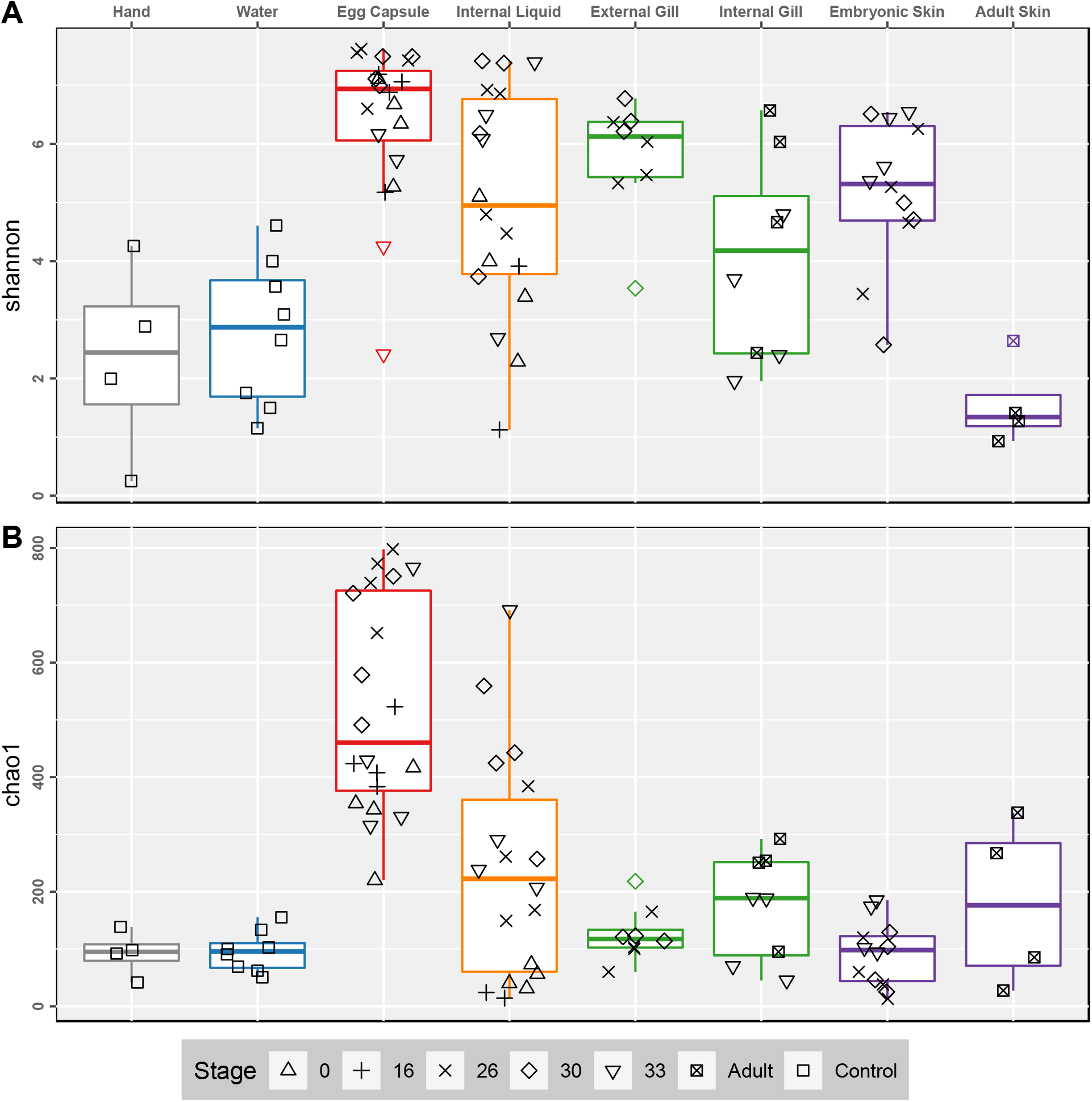
Alpha diversity of embryonic and adult skate bacterial communities. Boxplots of Shannon (A) and Chao1 (B) alpha diversity metrics for each sample site. Datapoints shown in black. Colored points are statistical outliers. Stage is indicated by shape. Grey: experimenters’ hands, blue: water, red: egg capsule, orange: internal liquid, green: gill, and purple: skin.

### Microbial source identification for each stage

We performed FEAST source tracking on each stage to assess the relative contribution of each sampling site in facilitating intergenerational transmission. FEAST estimates the relative proportional contribution of each potential source and assigns any unexplained components of the sink bacterial community to an unknown source [52]. A consequence of this emphasis on proportional mixing is limited power to identify the source of taxa with higher relative abundances in the sink compared to any source, leading to inflated unknown source values. Since some taxa are likely enriched during development of the little skate, FEAST values should be interpreted as identifying the most important sources at each stage, not directly estimating continuity between timepoints. Samples from the preceding embryonic stage were used as source pools for the target community (sink) of the focal stage to track microbial continuity until hatching. At stage 0, swabs from the representative adult females were used as the source pools (Fig. 4). For all timepoints, water samples were included as an environmental source pool, and the experimenters’ hands were also considered as a source of potential contamination.

**Figure 4:**
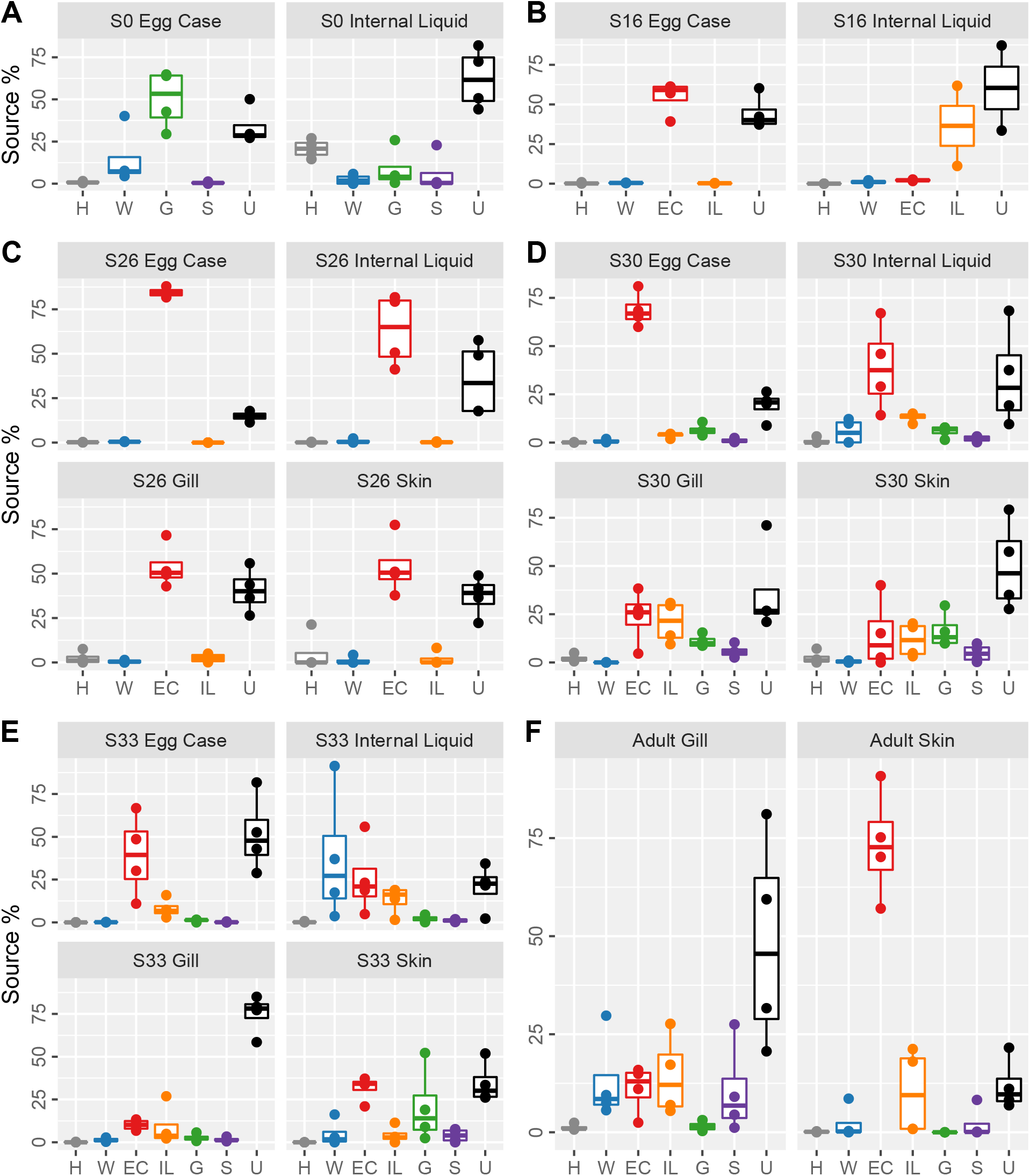
Source contributions to the little skate microbiota for each stage and tissue. Boxplots showing source contributions to the bacterial community of the skate microbiota at stage 0 (A), stage 16 (B), stage 26 (C), stage 30 (D), stage 33 (E) and adult (F) estimated using FEAST. Stage 0 used adult tissues as the source pools. Source contributions to adults (F) are shown for stage 33 source pools. Sources are colored as in Fig. 2. Letter codes refer to source pools from the previous stage or controls. H: Experimenters’ hands, W: water; EC: egg capsule, IL: egg capsule internal liquid, G: gill, S: skin, U: unknown source.

Hand samples are a negligible source for all samples except stage 0 internal liquid (hand contribution 20.1±5.5%, mean±standard deviation, Fig. 4A). Environmental water rarely contributes to the little skate microbiota; the exceptions are stage 0 egg capsule (14.8±16.9%) and stage 33 internal liquid (37.3±38.5%). In addition to water, the egg capsule microbiota at stage 0 is similar to the adult gill community (50.2±17.2%, Fig. 4A). At stage 16, the egg capsule and internal liquid microbiota are both largely conserved from the same tissue of the previous stage (54.6±10.4% and 36.5±35.7% continuity, respectively, Fig. 4B). At stage 26, egg capsule from the previous stage is the dominant bacterial source for all tissues (63.9±18.4%, Fig. 4C). At stage 30, egg capsule and internal liquid are sourced primarily from stage 26 egg capsule (68.6±8.9% and 39.0±22.7%, respectively), but gill and skin are sourced from a combination of the egg capsule, internal liquid, and gill of the previous stage (Fig. 4D). At stage 33, egg capsule remains contiguous (39.1±24.0%) while the internal liquid draws from egg capsule (25.6±21.6%) and internal liquid (13.2±8.2%) sources along with environmental water (Fig. 4E). The stage 33 gill microbiota, comprised of the earliest internal gill samples, is not particularly similar to any source while the skin is most similar to S30 egg capsule (31.6±7.2%).

To understand the extent to which taxa from the embryonic microbiota contributes to the adult bacterial communities, we ran FEAST on the adult samples using stage 33 tissues as the source pools (Fig. 4F, top). Adult gill has some contribution from stage 33 egg capsule (11.1±6.1%), internal liquid (14.3±10.3%), and skin (10.6±11.8%). The adult skin bacterial community is primarily derived from that of the late embryonic egg capsule (73.3±14.0%) (Fig. 4F).

### Identification of skate core microbiota

For subsequent analyses, we used Bray-Curtis dissimilarity to group samples of interest into four statistically significant and biological relevant groups: (1) egg capsule (all stages), (2) external gills (stages 26-30) and embryonic skin (stages 26-33) together, (3) internal gill (stage 33-adult), and (4) adult skin. Due to heterogeneity in the composition of the internal liquid throughout ontogeny and lack of large source contributions to other tissues, these samples are not considered in further analyses.

To extract the core microbiota of the little skate from our dataset, we identified taxonomic groups classified to the most specific level possible which were present in 75% or more of the samples comprising the four groups specified above (Supplementary Table S3). Egg capsule samples have a rich core microbial community with 212 identified taxonomic groups. Skate samples have smaller core microbiota: combined external gill and embryonic skin have ten, internal gills have twenty-two, and adult skin has fifty-six groups. There is high overlap between the core microbiota of these tissues (Table 1) and only egg capsule and adult skin house unique taxa (Supplementary Table S3).

**Table 1:** Core taxonomic units shared between tissues. List of taxa identified as part of the common core microbiota at a 75% presence cut-off in a least two of the following tissues: egg capsule, combined external gill (stages 26-30) and embryonic skin, internal gill (stage 33-adult), and adult skin.

| Taxa | Egg Capsule | External Gill/<br>Embryonic Skin | Internal Gill | Adult Skin |
| --- | --- | --- | --- | --- |
| *Actinomarinales* | x |  |  | x |
| *Alphaproteobacteria* | x | x | x | x |
| *Bacteroidia* | x | x |  | x |
| *Burkholderiaceae* | x |  |  | x |
| *Candidatus Nitrosopumilus* | x |  | x |  |
| *Chloroplast* | x |  | x | x |
| *Colwellia* | x |  | x | x |
| *Escherichia-Shigella* | x | x | x |  |
| *Flavobacteriaceae* | x | x | x | x |
| *Flavobacteriales* | x |  |  | x |
| *Gammaproteobacteria* | x | x | x | x |
| *Haliaceae* | x | x |  | x |
| *Lentisphaera* | x |  | x | x |
| *Pirellulaceae* | x | x | x | x |
| *Pseudoalteromonas* | x |  | x | x |
| *Pseudomonas* | x |  |  | x |
| *Rhizobiaceae* | x |  | x | x |
| *Rhodobacteraceae* | x | x | x | x |
| *Rhodothermaceae* | x |  |  | x |
| *Rubritalea* | x |  |  | x |
| *Saprospiraceae* | x | x | x | x |
| *SAR11 clade (Clade I)* | x |  | x | x |
| *Shewanella* | x |  | x |  |
| *Sphingomonadaceae* | x |  | x | x |
| *Sulfitobacter* | x |  |  | x |
| *Synechococcus CC9902* | x |  | x |  |
| *Thermoanaerobaculaceae*<br>Subgroup 10 | x |  |  | x |
| *Thiothrix* sp. FBR0112 | x |  |  | x |
| *Ulvibacter* | x |  | x |  |
| Uncultured verrucomicrobium<br>DEV007 | x |  | x |  |
| *Verrucomicrobium* sp. KLE1210 | x |  |  | x |
| *Vibrio* | x |  |  | x |
| *Vibrionaceae* | x |  | x | x |

We cross-referenced the genera identified as core to only the adult skin with the full taxa list for all samples in order to identify taxa exclusively present on adult tissues. This comparison identified six exclusively adult taxa. *Undibacterium* is the only genus found in all adult skin and gill samples that does not appear in any egg capsule or embryonic sample. *Spiroplasma, Salimicrobium*, and *Proprionivibrio* were each found in a single adult gill sample and in the adult skin core microbiota. *Sulfurospirillum sp. SM-5* and *Aeromonas* were unique to adult skin tissues.

### Differential abundance of bacterial taxa and predicted functions

For each of the four sample groups in Table 1, we used LEfSe analysis to identify taxa enriched in abundance (*P*<0.05, LDA>2; Supplementary Fig. 6). Adult skin is enriched in *Bacteroidia, Vibrio*, and *Mycoplasma agassizii*. Embryonic skin and the external gills are enriched in *Sphingomondales, Flavobacteriaceae*, and *Escheria-Shigella*. Internal gills are enriched in *Rhodobacterales* and *Alphaproteobacteria* of *SAR11 clade 1*. Finally, the egg capsule is characterized by higher abundance of *Alteromonadales, Pirellulales, Saprospiraceae*, and *Verrucomicrobia*.

Additionally, we used LEfSe to identify shifts in taxa abundance associated with the opening of the egg capsule slits. We compared open and closed samples of the egg capsule, internal liquid, and a combined set of the two tissues. While a few taxa were identified as statistically significant, closer inspection of abundances in each sample did not exhibit the expected pattern of similar levels of abundance across all samples in one condition compared to the other. Instead, significance was driven by differences in group means due to a few samples with high abundances in either the open or closed condition (Supplementary Fig. 7).

## Discussion

Our results show that the phyla *Proteobacteria* and *Bacteroidetes* comprise most of the bacterial communities associated with the little skate, as has been shown for other chondrichthyans [8, 9, 11–13]. Adult skate skin within our study has a uniquely high proportion of *Bacteroidetes* (>50%) compared to all other batoids [11, 12]. Within *Proteobacteria*, relative proportions of each class vary between chondrichthyan species, including the little skate [8, 9, 11–13]. Below the phylum-level, there is evidence of unique site-specific communities in our study. While all skate samples included *Gammaproteobacteria, Flavobacteriaceae, Pirellulaceae, Rhodobacteraceae*, and *Saprospiraceae*, we are unable to identify a common core microbial community below the family level for all skate samples across all developmental timepoints. Instead, our data suggest that early embryonic tissues support similar bacterial communities which differentiate into distinct internal gill and adult skin microbial communities later in development.

We sampled two parts of the egg capsule: the inner surface of the capsule and the internal liquid, which fills the space in the egg capsule not occupied by the developing embryo and yolk. The microbiota of the egg capsule has the highest taxonomic richness of any tissue sampled and has a complex core microbiota. This provides evidence that the egg capsule is a rich reservoir of bacteria for the developing embryo, similar to the dense microbial community observed in squid egg capsules [56, 57]. While the microbiota of the internal liquid is generally similar to egg capsule samples, there appears to be a collapse of *Actinobacteria* and *Bacilli* after stage 16, both rare in the egg capsule, with relative replacement by *Bacteriodia* and *Planctomycetacia*, both more abundant in the egg capsule. No taxa undergo consistent shifts in abundance in either the egg capsule or internal liquid upon opening.

Gills are the primary site of gas and waste exchange with the environment, offering a unique habitat for microbes. Little skate embryos develop transient external gill filaments between stages 25 and 32, which later regress into the body to form the adult internal gills [36]. During early stages, we find that the gill microbial community is undifferentiated from that of the embryonic skin. The mature gills, however, a harbor distinct microbial community, which is enriched for *Rhodobacterales* and *Alphaproteobacteria* of the *SAR11 clade*. These taxa are also enriched in the gills of reef fish, suggesting that marine vertebrate gills provide similar microbial environments [58]. Additionally, gills are the largest site of urea loss in skates [59]. This makes adult gill tissue particularly well-suited for a commensal relationship with *Nitrosopumilus*, which is known to use urea as an energy source [60], and is identified as a core bacterial genus in the gills starting at stage 33.

Adult skin has the lowest Shannon diversity of any tissue and unlike all other sites its microbiota is primarily composed of *Bacteroidetes* ASVs assigned to class *Bacteroidia* (81.2%). Previous work has shown that *Bacteroidetes* are dominant in many niches, are adapted to life on marine surfaces, play roles in polymer degradation, and contribute to immune function [61, 62]. Given the abundance of *Bacteroidetes* on skate skin, the functional implications of these ASVs on the host is a promising area for future research.

The low diversity observed in the adult skin samples may be due to the distinct properties of chondrichthyan skin, which is characterized by dermal denticles and a thin mucus layer [63]. Since the denticles do not develop until around the time of hatching, the biophysical properties of skate skin change after embryonic development is complete [64]. The skin is hypothesized to offer a selective microbial environment due to micropatterning of the dermal denticles, reduced laminar flow, and antimicrobial compounds [9, 65]. In support of this hypothesis, shark skin micropatterning has been shown to hinder microbial colonization and migration [66, 67]. Our data show that the adult skate skin bacterial community is distinct from that of embryonic skin, which clusters closely with the other embryonic tissues (Fig. 2). We hypothesize that the development of mature denticles shapes the bacterial community of adult chondrichthyan skin, though this hypothesis requires explicit testing. An alternate possibility for the low-diversity skin microbiota is that the taxa present possess antimicrobial properties and suppress competition. While stingray skin microbiota show some antibiotic potential, skate skin microbes exhibit lower levels of antibiotic activity [68, 69]. Future work is needed to assess the relative contributions of biophysical properties, antimicrobial secretions, and other mechanisms in shaping the bacterial community on chondrichthyan skin and the relevance of this community to host physiology and health.

Environmental sources likely contribute to the microbiota of adult skates. The one environmental source included in this study, the surrounding water, was not found to contribute meaningfully to gills or skin at any stage, however, deeper sampling may be necessary to detect low abundance taxa present in the water column. Identifying other potential environmental sources, such as diet and benthic substrates, for wild caught adults is not feasible and these sources could not be sampled for this study. Taxa identified only on adults cannot be explained by vertical transmission. In this study, we identified only six genera uniquely present on adult tissues. Of these six, only a single genus, *Undibacterium*, was present in all adult samples of both skin and gill. *Undibacterium* species have been isolated from other fishes and may play a role in biofilm degradation [70–73]. Given that this taxon was not detected in embryonic or water samples, skates likely acquire *Undibacterium* from an unknown environmental source, drastically enrich this genus from starting levels below the limit of detection, or through horizontal transmission.

We provide evidence of vertical transmission in an elasmobranch by tracking community continuity between different developmental stages and tissues. First, we found minimal water contributions to each sample, with the exception of internal liquid at stage 0 and after egg capsule opening. Second, skate samples have largely similar taxa between consecutive stages, with the largest shifts in community composition matching developmental changes in the physical properties of host tissues. Given the sealed nature of the egg capsule, differences in community composition are hypothesized to reflect enrichment of existing bacteria rather than recruitment from unknown sources, but this requires additional study to confirm. Finally, only six taxa were exclusively found on adult tissues, suggesting environmental sourcing may be limited. While these points are insufficient to show vertical transmission, taken together, along with the life history of the little skate, they imply vertical microbial transmission occurs. If future studies identify vertical transmission in this and additional chondrichthyan species, particularly those with alternate life histories, this would provide a potential mechanism underlying the signature of phylosymbiosis observed by others in elasmobranchs [11].

There are additional considerations for this dataset given our experimental design. Since individual egg capsules were not associated with particular adult females, direct pairwise comparisons between parent and offspring were not possible. Furthermore, embryonic sampling is lethal, so each developmental timepoint is comprised of different, unrelated embryos. Thus, inter-individual variation limits our ability to accurately track all ASVs between timepoints. While we did sample adult female skin at the cloaca, these samples are unlikely to capture the extent of diversity housed in the reproductive tract where the egg capsules form [25]. Like most elasmobranchs, little skates are polyandrous and multiple paternity is likely [74]. Similar promiscuity has been associated with higher microbial diversity in the female reproductive tract in other vertebrate groups [75–77], a pattern that may hold for skates. We hypothesize that the microbiota of the reproductive tract is highly diverse, potentially providing a richer source of microbiota to the egg capsule than is captured by the adult tissues sampled in this study. Nonetheless, the precise mechanisms by which the egg capsule is seeded with its rich bacterial community, and how transmission of pathogenic bacteria is minimized, await further investigation.

## Conclusions

This study provides the first exploration of the bacterial communities associated with the little skate throughout ontogeny and offers many intriguing possibilities for future microbiome research using this model chondrichthyan. Specifically, we identified a site-specific microbiota that is likely transferred between generations via the internal surface of the egg capsule and provide the first evidence that vertical transmission is present in an oviparous elasmobranch species.

## Supporting information

Supplementary Table 1: Amplification

Supplementary Table 2: Samples

Supplementary Table 3: Core Microbiota

## List of abbreviations

MRC: Marine Resources Center at the Marine Biological Laboratory, Woods Hole, MA, USA
ASV: Amplicon sequencing variant
FDR: False-discovery rate
LDA: Linear discriminant analysis
PC: Principal coordinate

## DECLARATIONS

### Ethics Approval

All procedures were conducted in accordance with Marine Biological Laboratory IACUC protocol 19-42.

### Consent for publication

Not applicable.

### Availability of data and materials

Raw sequencing data is available at the NCBI Short Read Archive (PRJNA688288). All scripts are available online at https://github.com/kmmika/Intergenerational-microbial-transmission-in-the-little-skate-Leucoraja-erinacea.

## Author Contributions

K.M. and A.S.O. conceived of and designed the project in consultation with N.H.S and D.M.W. K.M. and A.S.O. collected the data, performed the computational analyses, and wrote the manuscript. All authors edited and approved the final paper.

## Acknowledgements

The authors thank David Remsen, Dan Calzarette, and the MRC staff for assistance sampling the skates; Andrew Gillis for inspiration and staging the skate embryos, Tetsuya Nakamura for assistance with our IACUC protocol, Emily Davenport for directing us to computational resources, and Nipam H. Patel for use of lab space. We are deeply grateful to Tom A. Stewart, Terence D. Capellini, and the Carmody Lab for helpful comments on the manuscript.

## FIGURE AND TABLE LEGEND

**Supplementary Figure 1:**
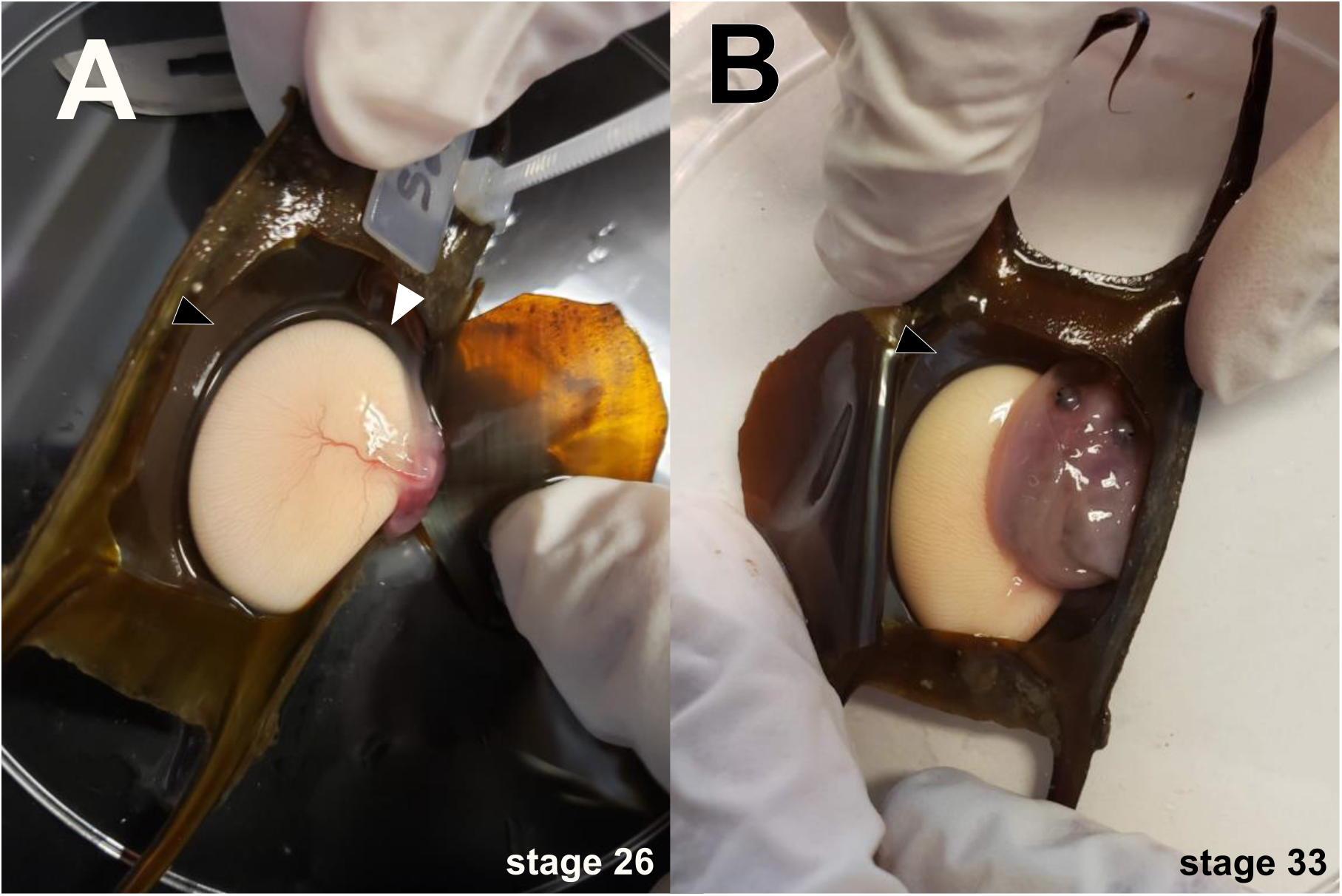
Internal liquid and egg capsule sampling locations. Egg capsules were windowed to access the developing embryo. Black arrowheads indicate the internal surface of the egg capsule. White arrowhead in A indicates where the transparent, gelatinous internal liquid of closed egg capsules was collected. Open egg capsules were directly drained into microcentrifuge tubes before windowing.

**Supplementary Figure 2:**
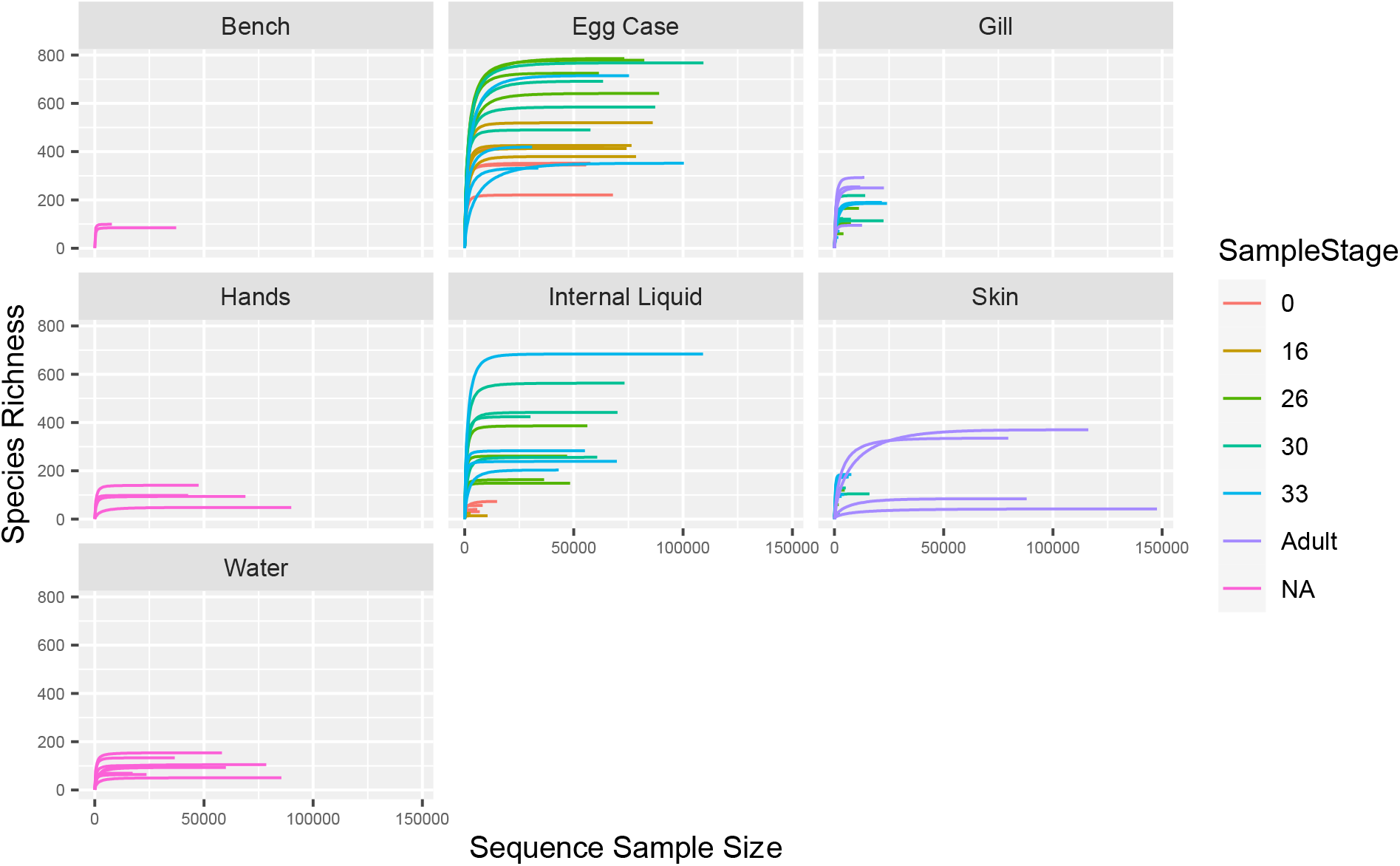
ASV rarefaction curves of samples. While number of reads varied between samples, species richness plateaus for each below the maximum sequencing depth.

**Supplementary Figure 3:**
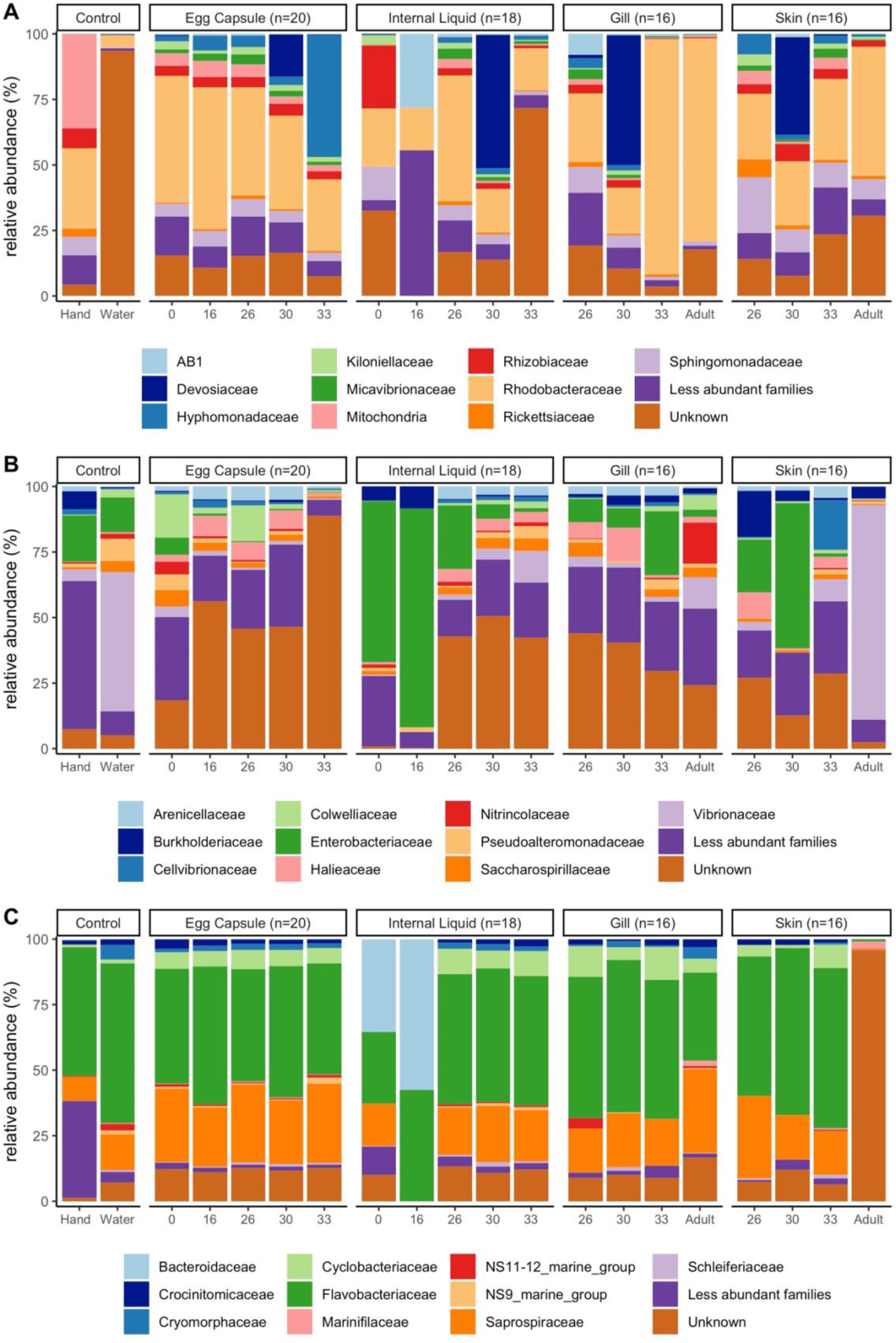
Family-level composition of embryonic and adult skate bacterial communities. Relative abundance of the top ten bacterial families in the classes *Alphaproteobacteria* (A), *Gammaproteobacteria* (B), and *Bacteroidia* (C) in the dataset are shown for each site and timepoint as well as for water and hand controls. For the controls, n=4 for hand and n=8 for water samples.

**Supplementary Figure 4:**
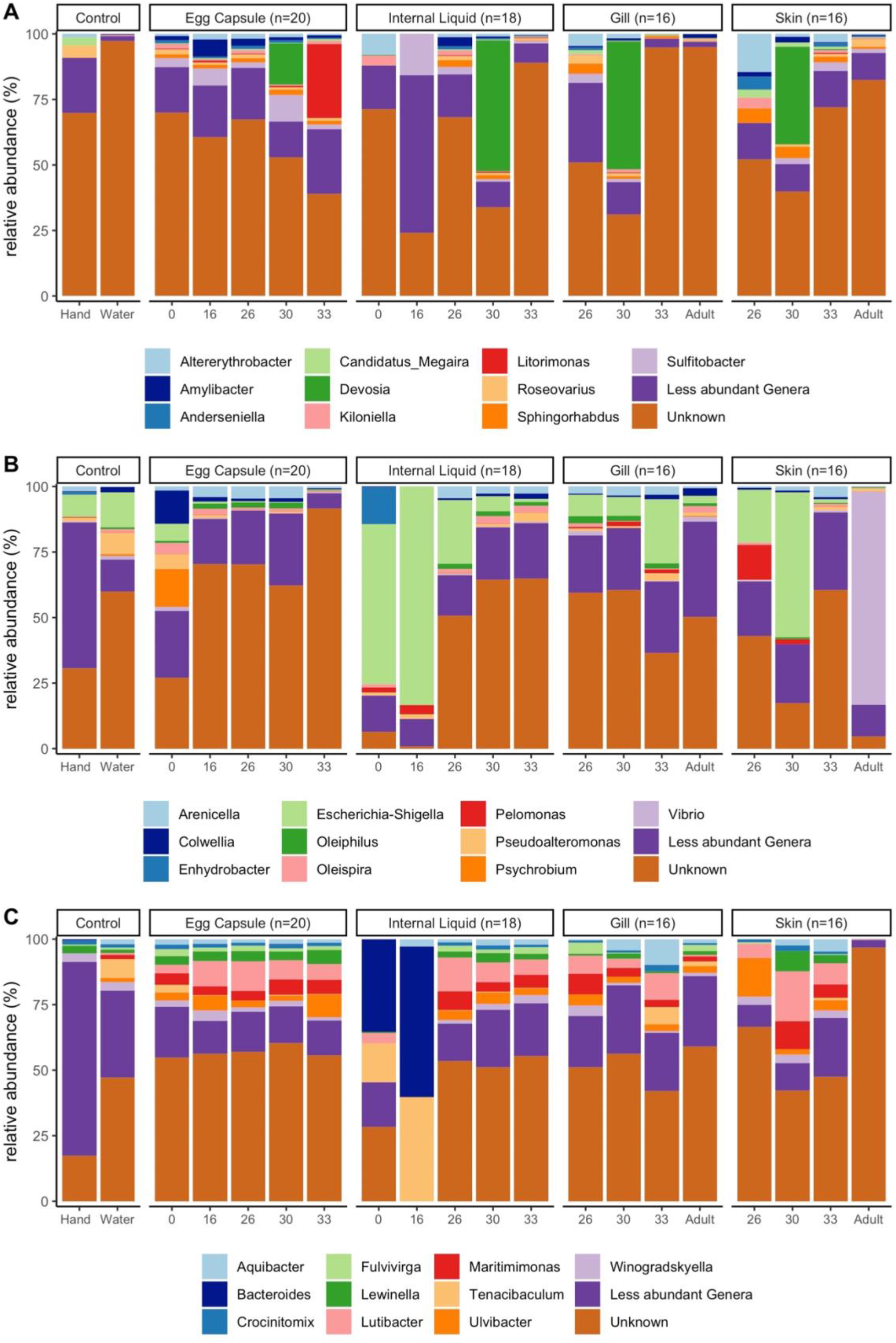
Genus-level composition of embryonic and adult skate bacterial communities. Relative abundance of the top ten bacterial genera in the classes *Alphaproteobacteria* (A), *Gammaproteobacteria* (B), and *Bacteroidia* (C) in the dataset are shown for each site and timepoint as well as for water and hand controls. For the controls, n=4 for hand and n=8 for water samples.

**Supplementary Figure 5:**
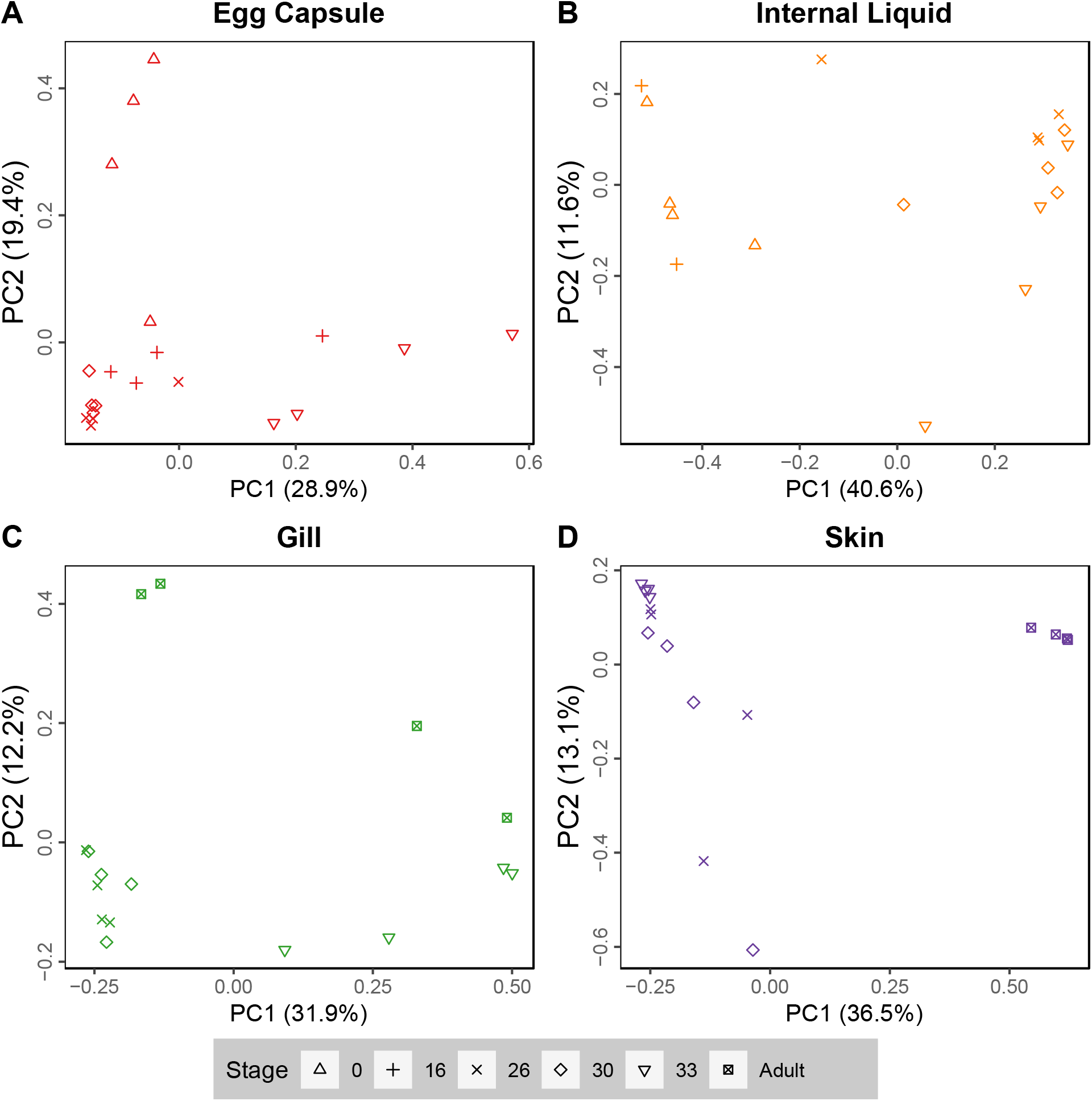
Principal coordinate analysis plots of bacterial communities by tissue. PCoA analysis (Bray-Curtis) plots of PC1 versus PC2 for (A) egg capsule, (B) Internal liquid, (C) gill, and (D) skin samples.

**Supplementary Figure 6:**
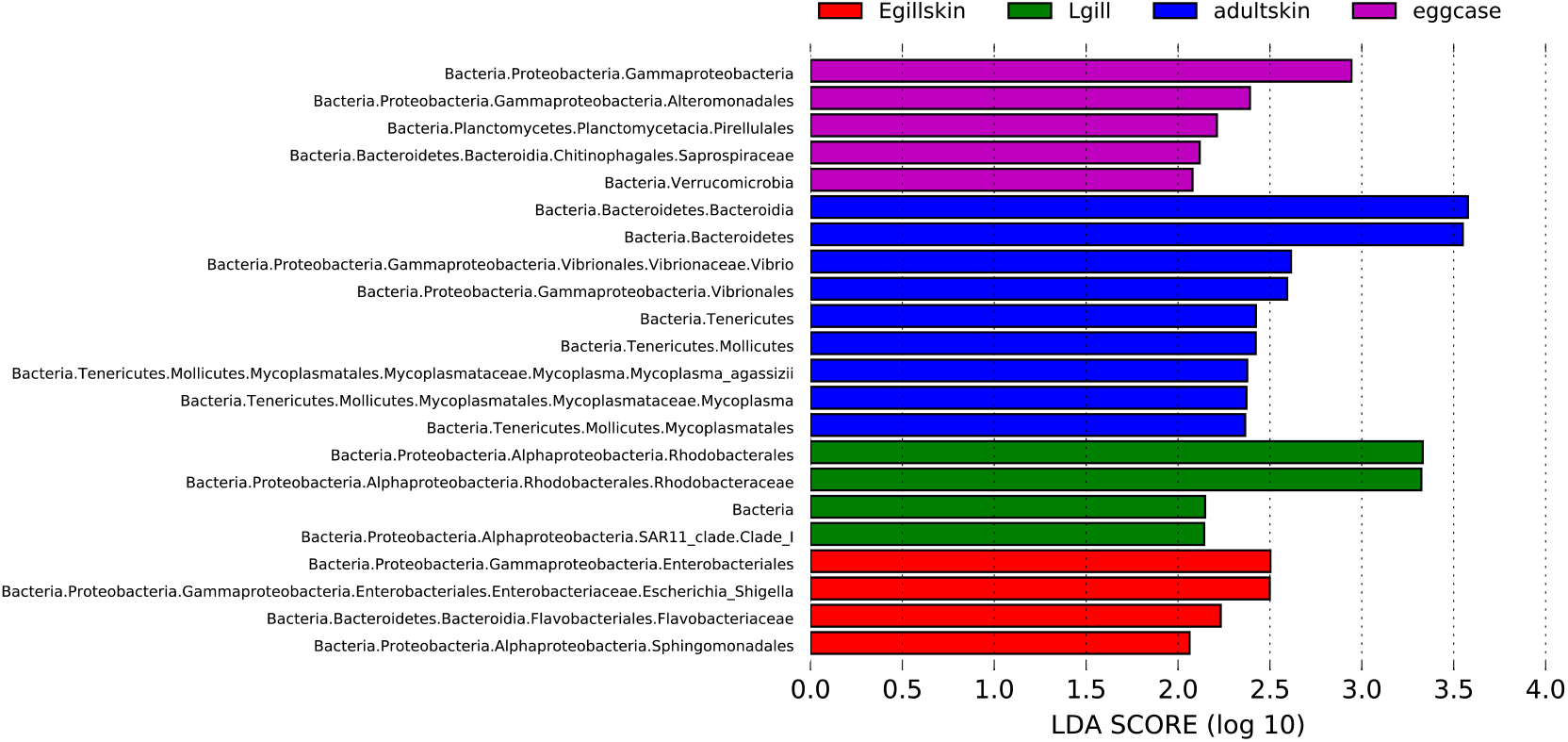
Differentially abundant bacterial taxa among skate tissues. LEfSe analysis at *P* <0.05 and LDA>2. Only differentially abundant tree branches are shown. eggcase: egg capsule, Egillskin: embryonic external gills (stages 16–30) and embryonic skin (stages 16–33), Lgill: internal gill (stage 33-Adult), adultskin: adult skin.

**Supplementary Figure 7:**
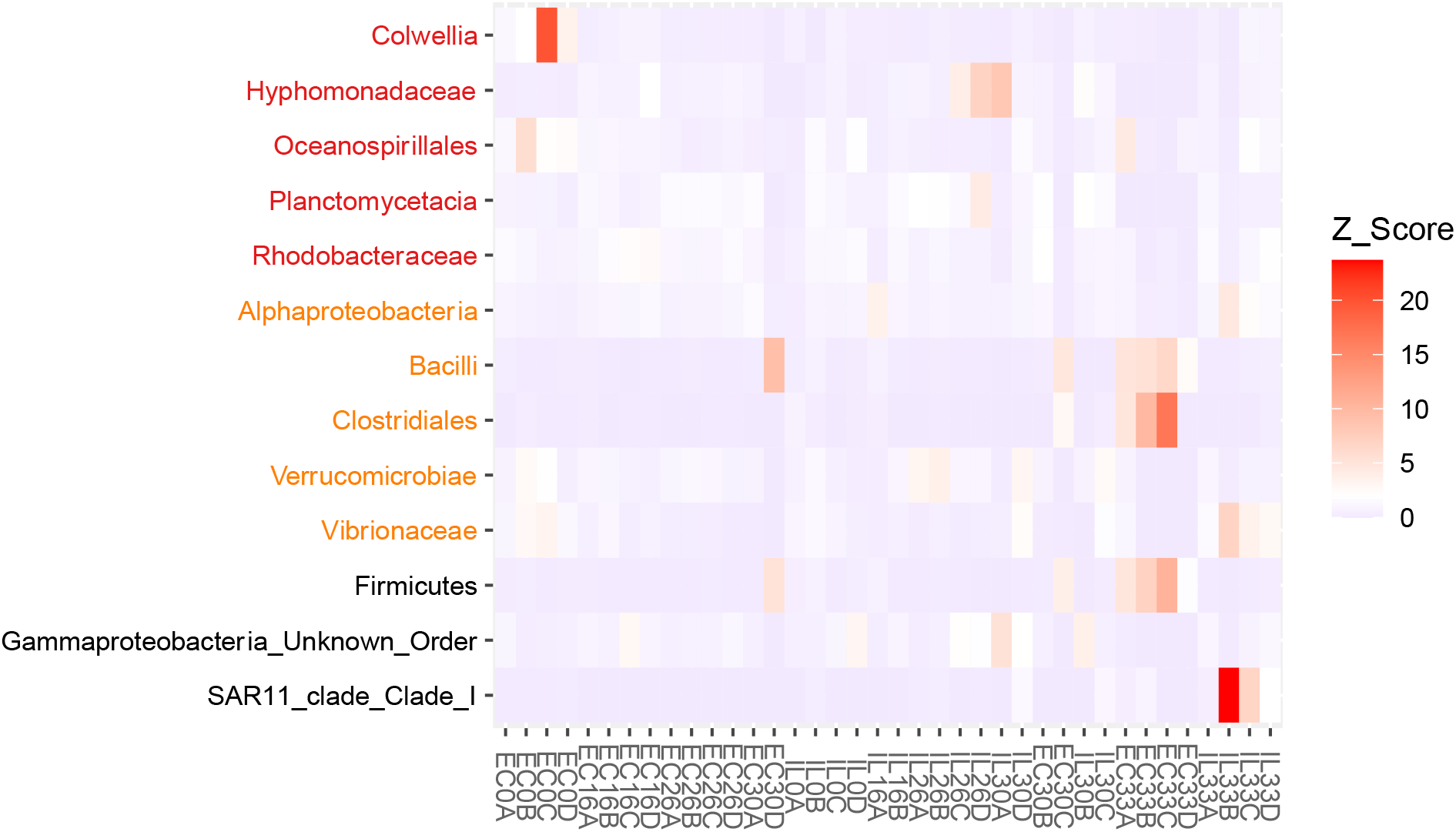
Significant taxa comparisons in closed versus open egg capsules. Heatmap of the taxa identified by LEfSe (*P*<0.05, LDA>2) at the lowest categorized taxonomic level for comparisons of open versus closed egg capsule (red), internal liquid (orange) and combined egg capsule and internal liquid (black). Abundances of each sample are shown as Z-scores. Open samples include stage 30 replicates A & D, as well as all stage 33 samples. All other samples come from closed egg capsules.

**Supplementary Table 1: Sample amplification results**

<u>Table S1 Amplification.xlsx</u>

**Supplementary Table 2: Sample reads summary statistics**

Table S2 Samples.xlsx

**Supplementary Table 3: Complete core microbiota results**

Table S3 Core Microbiota.xlsx

